# Cancer Drug Bortezomib, a Proteasomal Inhibitor, Triggers Cytotoxicity in Microvascular Endothelial Cells via Multi-Organelle Stress

**DOI:** 10.1101/2025.03.25.645304

**Authors:** Prajakta Sawant, Aleena Mathew, Johanna Bensalel, Julio Gallego-Delgado, Pratyusha Mandal

## Abstract

Proteasomes maintain cellular homeostasis by degrading abnormal proteins, while cancer cells exploit them for survival, becoming a key chemotherapeutic target. Bortezomib (BTZ), a reversible proteasomal inhibitor, is a front-line treatment for multiple myeloma, mantle cell lymphoma, and non-small cell lung cancer. However, its efficacy is limited by severe side effects, including neurotoxicity and cardiovascular distress, with its toxicity mechanisms largely unexplored. Here, we discover that Bortezomib (BTZ), is cytotoxic to non-cancerous cells distinctly from Carfilzomib (CFZ), the second-line irreversible PI. BTZ or CFZ is administered intravenously, impacting blood vessel (vascular) endothelial cells. We used human pulmonary microvascular endothelial cells (HPMECs) to demonstrate that BTZ but not CFZ elicits endoplasmic reticulum (ER) stress, mitochondrial membrane compromise, mitochondrial reactive oxygen species (ROS) accumulation, and Caspase (CASP)9 activation (mediator of Intrinsic apoptosis) within fifteen hours of treatment. By twenty-four hours, BTZ-treated cells display cleavage of CASP8 (mediator of extrinsic apoptosis), activation of CASP3 (terminal executioner of apoptosis), cell-death and vascular barrier loss. Pan-caspase inhibitor zVAD significantly rescues BTZ-treated cells from cytotoxicity. Both BTZ and CFZ effectively kill MM cells. These findings reveal novel insights into fundamental signaling of regular cells where reversible inhibition of the proteasome dictates a unique cascade of stress distinct from irreversible inhibition. These harmful effects of BTZ emphasize the need to re- evaluate its use as a frontline chemotherapy for MM.

**Graphical Abstract:** 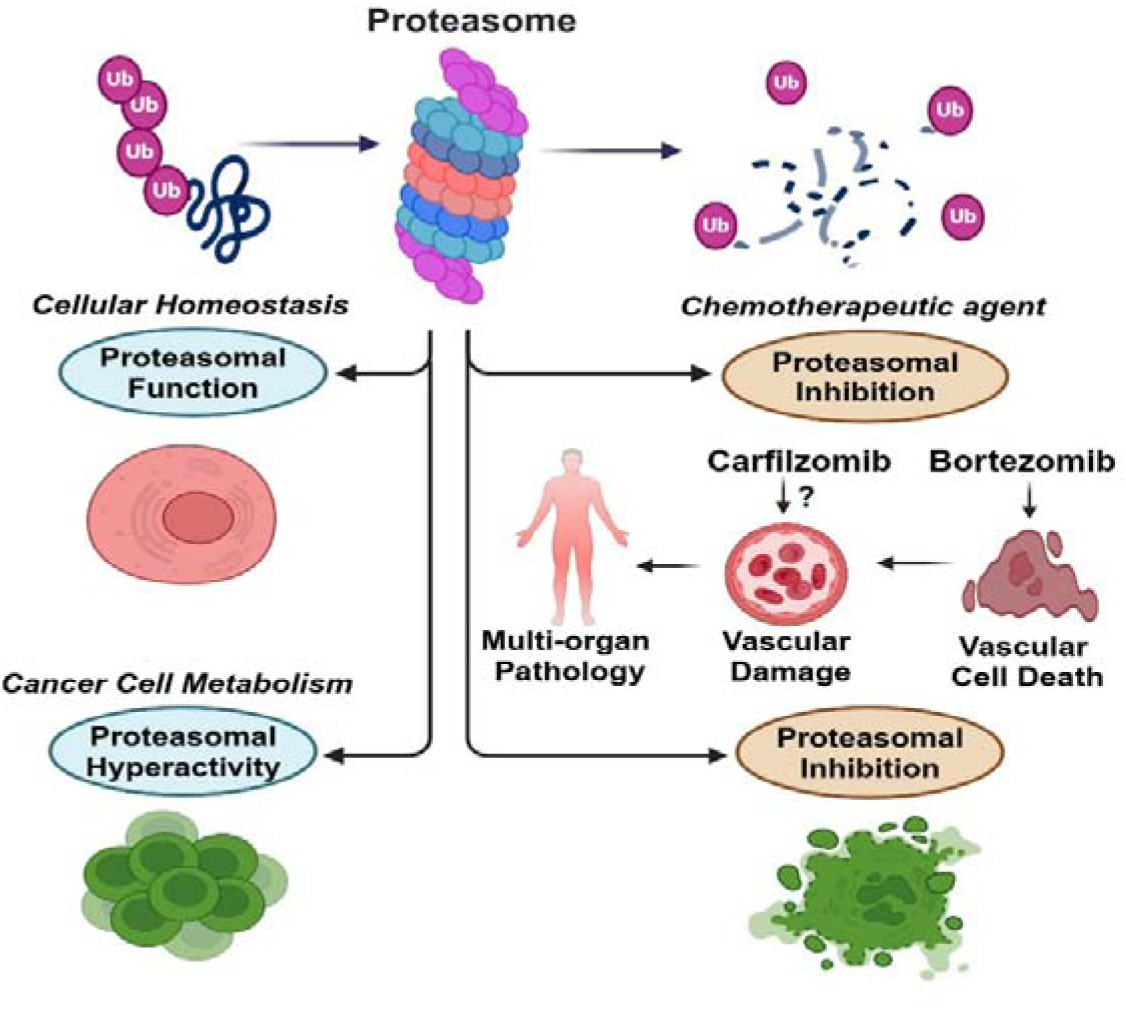

**Highlights:**

- Reversible proteasomal inhibitor Bortezomib is cytotoxic to non-cancerous, microvascular endothelial cells
- In endothelial cells, Bortezomib, but not irreversible inhibitor Carfilzomib, activates temporal cascade of caspases (Caspase-9, Caspase-8, Caspase-3) triggering apoptosis
- Caspase activation results from ER stress (via the IRE1α-CHOP) pathway and mitochondrial stress (ROS accumulation) independently from contribution from extrinsic signal via TNF
- Bortezomib-dependent cytotoxicity compromises endothelial barrier potential

## Introduction

Proteasomes are intricate protein complexes that play a crucial role in maintaining cellular homeostasis by regulating protein turnover, degrading misfolded or unwanted proteins, and generating peptides essential for immune responses. The mammalian 26S proteasome consists of two subcomplexes: the 20S catalytic core and the 19S regulatory particle, which caps both ends of the core [1]. In cancer cells, the upregulation of proteasome activity is vital for survival, as these cells produce large quantities of proteins and rely on efficient degradation of damaged proteins involved in cell cycle progression and apoptosis [2].

Building on this, proteasomal inhibitors (PIs) have become a key class of chemotherapeutic agents, with particular utility in treating multiple myeloma (MM) [3]. The first-in-class proteasome inhibitor, Bortezomib (BTZ), was approved in 2003 for relapsed and refractory MM, becoming the first-line therapy in 2008. It is also approved for mantle cell lymphoma and has shown promise in preclinical studies for treating solid tumors like non-small-cell lung cancer (NSCLC) [4]. However, despite its clinical efficacy, BTZ is associated with severe pulmonary complications, renal toxicity, and peripheral neuropathy [5–11]. Here, we demonstrate that BTZ induces cytotoxicity in human pulmonary microvascular endothelial cells (HPMECs) via activation of intrinsic cellular stress pathways, whereas Carfilzomib (CFZ), an irreversible proteasomal inhibitor, does not elicit similar cytotoxic effects, despite both drugs effectively sensitizing MM cells to cell death.

BTZ is a reversible inhibitor that binds to the catalytic β5/chymotrypsin site of the 26S proteasome [12]. Inhibition occurs within one hour of treatment, and activity typically returns to normal within 72 to 96 hours post-administration [13]. CFZ, approved in 2012 for advanced MM monotherapy and in 2015 for combination therapies, irreversibly binds to the N-terminal threonine-containing active sites of the 20S proteasome [14]. Both inhibitors lead to protein accumulation, endoplasmic reticulum (ER) stress, mitochondrial stress, reactive oxygen species (ROS) production, unfolded protein response (UPR), cell cycle arrest, and apoptosis in cancer cells [15–17]. Apoptosis can occur via two primary pathways: intrinsic apoptosis, triggered by mitochondrial stress and regulated by BCL2 family proteins [18], and extrinsic apoptosis, which involves caspase 8 activation following death receptor signaling [19]. Under normal conditions, intrinsic apoptosis is inhibited by anti-apoptotic proteins such as BCL2, BCLXL, and MCL1. Inhibition of these proteins promotes the activation of BAX and BAK, which oligomerize on the mitochondrial membrane, compromising membrane integrity and causing cytochrome c release [20]. This results in CASP9 activation, which subsequently activates CASP3, leading to cell death. BTZ treatment reduces BCL2 expression, triggering intrinsic apoptosis in MM cells [21, 22]. BTZ also activates extrinsic apoptosis programs, including stress-responsive autophagy [23–24]. Crosstalk between intrinsic and extrinsic apoptosis occurs when CASP8 cleaves Bid into its truncated form (tBid), which disrupts mitochondrial integrity and feeds into CASP9-dependent CASP3 activation [25]. Like BTZ, CFZ induces intrinsic and extrinsic apoptosis as well as autophagy in cancer cells [14, 26–30]. Both drugs also drive mitochondrial ROS production in MM cells [31]. ROS can activate JNK and p38 MAPK pathways, further promoting intrinsic apoptosis [32, 33]. These observations underscore the importance of ER stress, mitochondrial stress, and UPR in sensitizing cancer cells to proteasomal inhibitors. However, despite these insights into PI effects on cancer cells, less is understood about the impacts of mode of proteasomal inhibition on non-cancerous cells particularly concerning the adverse effects observed in patients. Neurotoxicity is the primary side effect seen with both BTZ and CFZ treatment, with BTZ specifically inducing higher proteotoxic stress on human neurons [11]. Clinically, BTZ is administered intravenously or subcutaneously, while CFZ is typically administered intravenously [34, 35]. Thus, there is strong potential for interaction with blood vessel endothelial cells. Vascular damage is a noted side effect of PIs [36]. To address this deficit in knowledge, in this study, we focus on the effect of reversible (BTZ) and irreversible (CFZ) proteasomal inhibitors on human pulmonary microvascular endothelial cells (HPMECs), as pulmonary toxicity is a known critical adverse effect of BTZ [37–39]. We demonstrate that BTZ induces ER stress, mitochondrial membrane disruption, ROS accumulation, and CASP9 activation in HPMECs within 15 hours of treatment. By 24 hours, BTZ-treated cells show CASP8 cleavage without detectable TNF, suggesting that CASP8 activation is primarily due to intrinsic cellular stress rather than cell-extrinsic signaling. These cellular stress responses ultimately lead to cell death and loss of cell-cell integrity. In contrast, CFZ does not induce similar cytotoxic effects in HPMECs. While both BTZ and CFZ effectively induce cell death in MM cells (MM1.S). These results provide novel insights into how the mode of proteasomal inhibition determines the fate in cancerous and non-cancerous cells. Our findings also raise questions about the use of BTZ as a frontline chemotherapy agent, particularly in light of its off-target effects on endothelial cells and potential for severe pulmonary toxicity. The affordability and widespread availability of BTZ make it a cost-effective treatment for MM, but its side effects disproportionately affect low-income populations, exacerbating racial and socioeconomic disparities in access to safer treatments [40]. Our study underscores the need for a) re-evaluating the use of BTZ in chemotherapy and the development of more targeted drugs to minimize off-target effects and improve patient outcomes; b) consider application of BTZ in combination with therapies that promote vascular health. Additionally, our work provides crucial information for developing strategies to mitigate the pulmonary and other toxicities associated with BTZ, ultimately contributing to the safer use of proteasomal inhibitors in clinical settings.

## Results

### Reversible Versus Irreversible Proteasomal Inhibition Drives Different Outcomes in Cancer and Non-Cancer Cells

We chose 100 nM of drug concentration based on prior studies in cancer cells to evaluate reversible and irreversible proteasomal inhibitors. At this dose, BTZ or CFZ fully blocks proteasomal function and induces death in multiple myeloma cells [41]. Similar concentrations have been used to assess the BTZ’s *in-vivo* toxicity [11]. When we treated MM1.S cells, with 100 nM of CFZ or BTZ, both drugs killed cells compared to vehicle control DMSO within 12 hours of treatment based on loss of ATP (Figure 1A, left panel). For HPMECs, 24 hours of treatment with BTZ resulted in a significant loss of ATP in comparison to DMSO (Figure 1A right panel). In contrast, CFZ-treated cells did not die. We confirmed the loss of viability using bright-field (Figure 1B) and fluorescence (Supplementary Figure 1A) microscopy. Morphological analysis revealed that BTZ treatment led to the loss of adherent cells, resulting in floating circular cellular bodies (Figure 1B). Cells also displayed punctate green, fluorescent signals of SYTOX incorporation (Supplementary Figure 1A).

**Fig 1.**
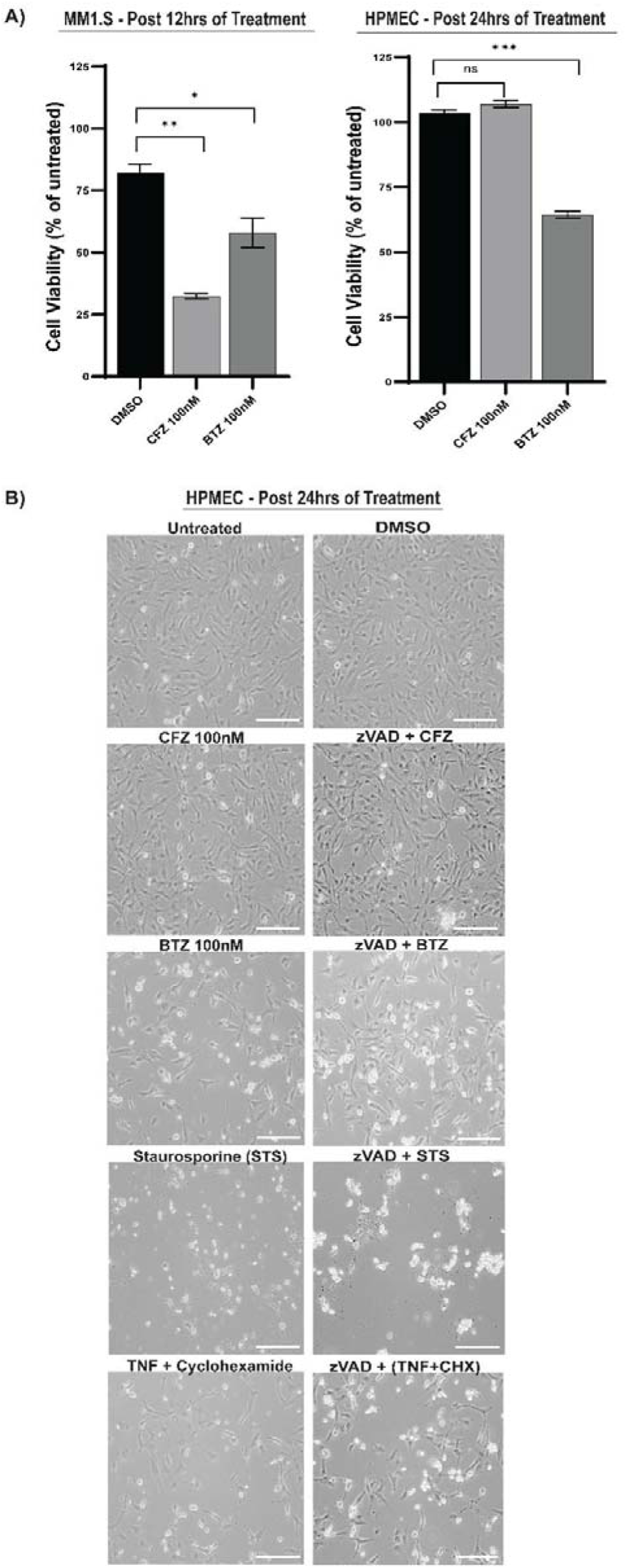
Bortezomib (BTZ) and Carfilzomib (CFZ) response in multiple myeloma (MM1.S) cells versus human pulmonary endothelial cells (HPMEC). (A) Cell viability measured by ATP levels in MM1.S (left) and HPMECs (right) after 100 nM CFZ or BTZ treatment compared to DMSO control. Data are presented as viability percentage relative to untreated cells (mean ± SEM, n = 6–12). Statistical significance assessed by Mann-Whitney test: *p < 0.01 CFZ 100 nM vs. DMSO, **p = 0.0001 BTZ 100 nM vs. DMSO (right); no significant difference for CFZ 100 nM vs. DMSO (left). (B) Bright-field microscopy images of HPMECs after 24-hour treatment with DMSO (1%), CFZ (100 nM), BTZ (100 nM), STS (1 μM), TNF (25 ng/ml) + Cycloheximide (CHX 1 μM), or zVAD (20-25 μM) in combination with CFZ, BTZ, STS, or TNF+CHX. Scale bar = 170 μm. Representative images from three independent experiments (n = 3 biological replicates).

SYTOX, a cell-impermeable fluorescent dye, exhibits a punctate pattern upon membrane permeability, a hallmark of dying cells [42]. More CFZ-treated cells appeared to remain viable. These observations stand in contrast with previous work showing that 100 nM of BTZ or CFZ does not cause death in human neurons [43], suggesting that the observed toxicity is cell-type dependent. To define how HPMECs die, we treated cells with BTZ, CFZ, vehicle control, or established death controls. Controls included staurosporine (STS), an inducer of mitochondria-mediated cell death, including intrinsic apoptosis [44], and TNF plus cycloheximide, which activates CASP8-dependent extrinsic apoptosis. [45] (Figure 1B, Supplementary Figure 1A). Bright-field microscopy (Figure 1B) revealed cell loss and increased punctate SYTOX signaling following STS or TNF+CHX treatment, confirming HPMEC death with these stimuli. Pan-caspase inhibition (zVAD) did not alter STS- or TNF+CHX-induced phenotypes (Figure 1B, Supplementary Figure 1A). In BTZ-treated cells, zVAD partially rescued cell death and SYTOX inclusion. Additionally, no evidence of necroptosis was observed. Thus, non-cancerous HPMECs are susceptible to BTZ-induced, caspase-dependent death at 100 nM, but not CFZ.

### BTZ Triggers Apoptotic Signals in MM1.S Cells and HPEMCs

We treated MM1.S and HPMECs with CFZ or BTZ and analysed cytoplasmic proteins by immunoblotting (Figure 2, Supplementary Figure 2). MM1.S cells showed apoptotic markers cleaved CASP3 and Cl-PARP (Figure 2A, Supplementary Figure 2A). STS induced these markers, confirming its role in intrinsic apoptosis. zVAD reduced both Cl-CASP3 and Cl-PARP in MM1.S cells, indicating caspase-dependent cascade (Figure 2A, Supplementary 2A). CFZ, BTZ, and STS induced Cl-PARP in HPMECs (Figure 2A, Supplementary Figure 2A). Cl-CASP3 appeared only in STS-treated control cells (Figure 2A, Supplementary Figure 2A). Neither CFZ nor BTZ induced Cl-CASP3 in HPMECs at 15 hours, indicating cell-type-specific variability in proteasomal inhibition resistance. zVAD reduced cleaved Cl-CASP3 in STS-treated HPMECs (Supplementary Figure 2A). These data show similar caspase-dependent kinetics in myeloma and endothelial cells, though proteasome inhibitor effects differ. To explore upstream signaling, we assessed Cl-CASP9. CASP9 is a crucial initiator of intrinsic apoptosis, activated by cell stress and proteasomal inhibition in cancer cells. STS treatment induced CASP9 cleavage, producing expected 35 kDa fragments, along with a smaller ∼20 kDa and larger ∼90 kDa products (Figure 2B, Supplementary 2B). CFZ didn’t induce CASP9 cleavage (Figure 2B, Supplementary 2B), but BTZ induced both the cleaved 35 kDa CASP9 product and the ∼20 kDa form (Figure 2B, Supplementary 2B). BTZ-treated HPMECs activated CASP9 without executing apoptosis at 15 hours, suggesting stress signaling activation. zVAD increased the 35 kDa cleaved CASP9 form (Figure 2B, Supplementary 2B), revealing new insights into CASP9 regulation. While zVAD inhibits CASP9 activity, it does not prevent all cleavage products. CFZ plus zVAD-treated cells, with accumulated Cl-CASP9, showed no contribution to cell death. Studies suggest zVAD partially inhibits CASP9 autocleavage by inducing conformational changes that prevent the formation of mature CASP9 [46,47] (Figure 2B, Supplementary 2B). Caspase activation may inhibit proteasome function, enhancing proteasomal effects, and explaining Cl-CASP9 appearance [48]. Overall, BTZ induces intrinsic apoptotic signaling in HPMECs. Further, we evaluated extrinsic apoptotic signaling via Cl-CASP8. At 15 hours, Cl-CASP8 appeared in STS-treated cells only. (two expected products ∼43 kDa and ∼18 kDa) in (Figure 2C, Supplementary Figure 2C). By 24 hours, STS-induced CASP8 cleavage was no longer evident, likely due to cellular content loss, indicating a temporal CASP8 activation. TNF + CHX did not cleave CASP8, suggesting differential TNF signaling in HPMECs. In contrast, BTZ treatment induced Cl-CASP8 at 24 hours, with zVAD rescuing this activation (Figure 2C, Supplementary 2C), a phenomenon not seen at 15 hours. At 24 hours, BTZ-treated cells also showed elevated Cl-CASP3 and Cl-PARP levels (Figure 2D), indicating a unique caspase cascade in HPMECs (CASP9→CASP8→CASP3). Both BTZ and STS treatments caused Cl-CASP3 laddering, while TNF + CHX treatment also elevated Cl-CASP3. zVAD altered Cl-CASP3 patterns and partially rescued cell death, suggesting regulatory effects on caspase activation. BTZ plus zVAD partially rescued death, with apoptosis driven by Cl- CASP3 fragments. These findings reveal a caspase activation pattern in BTZ-induced cell death: CASP9→CASP8→CASP3, leading to both intrinsic and extrinsic apoptosis. LC3B levels in BTZ or CFZ-treated HPMECs showed no significant change, ruling out autophagy involvement (Supplementary Figure 2C). No evidence of necroptosis was found in any treatment.

**Fig. 2.**
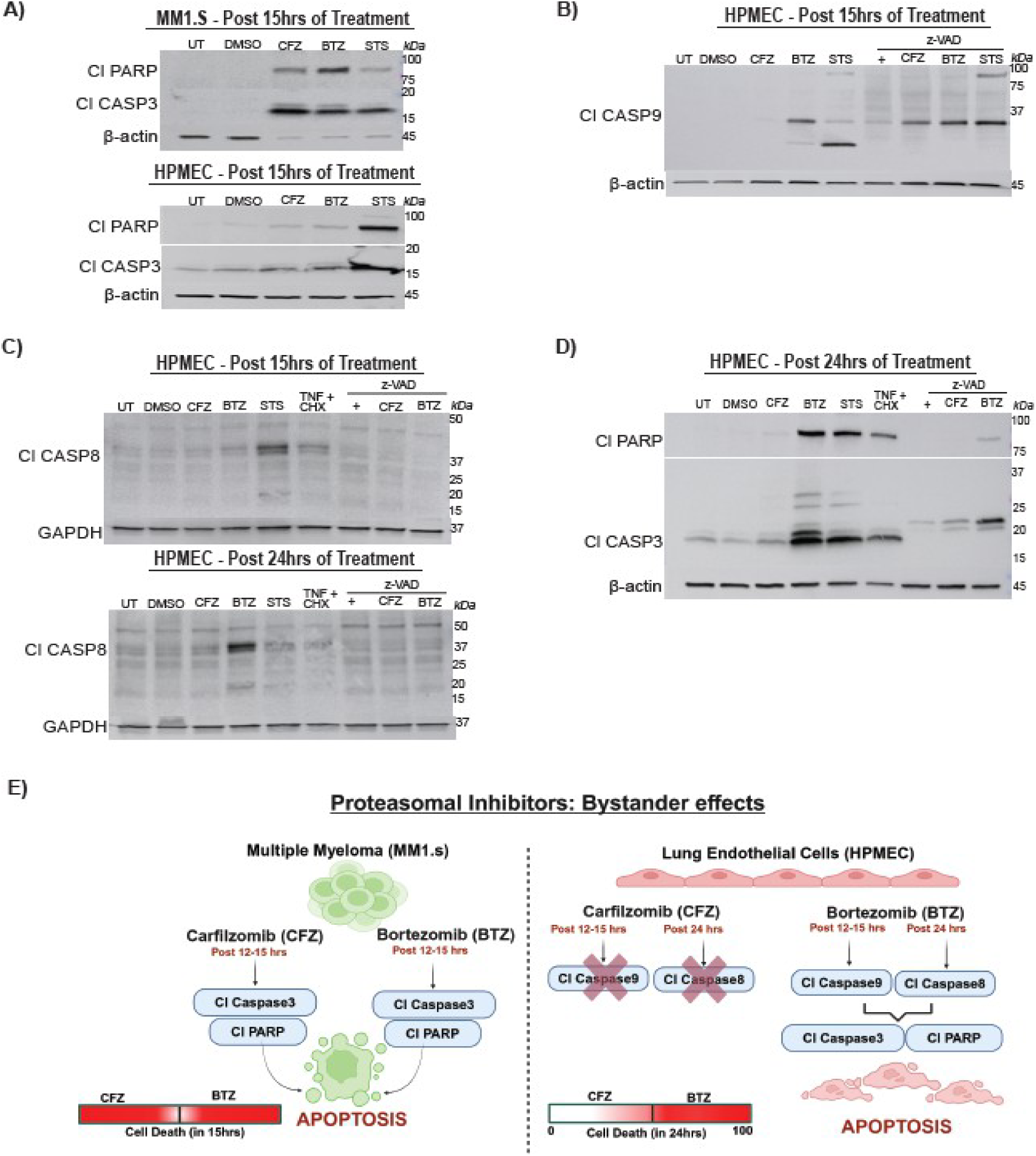
Bortezomib induces apoptosis in pulmonary endothelial cells. (A-D) Representative immunoblot analyses of cell death markers at the indicated time points following treatment with DMSO, CFZ (100 nM), BTZ (100 nM), STS (1 μM), TNF (25 ng/ml) + CHX (1 μM), zVAD (20-25 μM), and combinations of zVAD with CFZ, BTZ, or STS. (A) MM1.S (upper) and HPMEC (lower) showing cleaved CASP3 (Cl-CASP3; 17, 19 kDa) and cleaved PARP (Cl-PARP; 89 kDa), with β-actin (45 kDa) as loading control. (B) HPMEC showing cleaved CASP9 (Cl-CASP9; 35 kDa), with β-actin (45 kDa) as control. (C) HPMEC showing cleaved CASP8 (Cl-CASP8; 43, 18 kDa), with GAPDH (37 kDa) as control. (D) HPMEC showing cleaved CASP3 (Cl-CASP3; 17, 19 kDa) and cleaved PARP (Cl-PARP; 89 kDa), with β-actin (45 kDa) as loading control. (E) Schematic illustration of apoptosis signaling pathways activated by proteasomal inhibitors in MM1.S and HPMEC cells.

### BTZ Drives ER Stress in HPMECs

We next investigated cell-intrinsic stress mechanisms mediating BTZ-induced intrinsic cell death. In cancer cells, PIs drive ER and mitochondrial stress, leading to intrinsic apoptosis [49], but their roles in non-cancer cells remain unclear. To address this, we analysed ER stress markers via immunoblotting (Figure 3A). We assessed ER stress markers at 15 hours post-treatment, coinciding with CASP9 activation, as ER stress is a known upstream driver of this pathway. CFZ treatment induced CHOP over DMSO-treated cells. BTZ induced a further elevation in levels of CHOP over CFZ (Figure 3A). CHOP is a key ER stress marker induced via PERK-, IRE1-, or ATF6-mediated pathways (Figure 3B). PERK, activated by unfolded proteins, phosphorylates eIF2α, leading to ATF4-driven *CHOP* expression [50]. In the IRE1 pathway, unfolded proteins trigger IRE1α autophosphorylation, activating XBP-1, which upregulates apoptotic genes, including *Chop* [51]. PERK was detected in unmanipulated and DMSO-treated control cells, with no significant elevation under CFZ or BTZ conditions (Figure 3A). IRE1α was detected in unmanipulated or DMSO-treated cells with CFZ increasing its expression. BTZ treatment further enhanced levels of IRE1α (Figure 3A). This shows that in HPMECs, reversible proteasomal inhibition by BTZ induced IRE1α➔CHOP➔ER stress pathway significantly more than CFZ. This also coincided with BTZ-dependent CASP9 activation (Figure 2B, Supplementary Figure 2B), CASP8 activation (Figure 2C lower panel) and cell death (Figure 1 and Supplementary Figure 1). As with CASP3 and CASP9, zVAD did not eliminate increased CHOP levels in BTZ-treated cells (Figure 3A). zVAD reduced IRE1α levels in all conditions (Figure 3A). Therefore, caspase activity regulates BTZ-driven ER stress. To determine whether CFZ and BTZ differ further upstream in the ER stress pathway, we analyzed BiP (Figure 3A), a chaperone protein that acts as a primary ER stress sensor by interacting with the luminal domains of UPR proteins IRE1α or PERK [52]. DMSO or CFZ or BTZ all induced comparable BiP expression compared to unmanipulated cells (Figure 3A). BiP appeared as a doublet, suggesting a DMSO-induced ER response with no additional effect from CFZ or BTZ. zVAD, with or without drug treatment, further enhanced the doublet, highlighting caspase-dependent regulation of ER stress in HPMECs. Caspase inhibition likely leads to protein accumulation in the cytoplasm. In conclusion, we show that BTZ induces ER stress in normal cells as seen in cancer cells [49]. BTZ but not CFZ, specifically induces the IRE1α-CHOP axis, rather than a generalized UPR activation without altering protein chaperoning inside the ER.

**Fig. 3.**
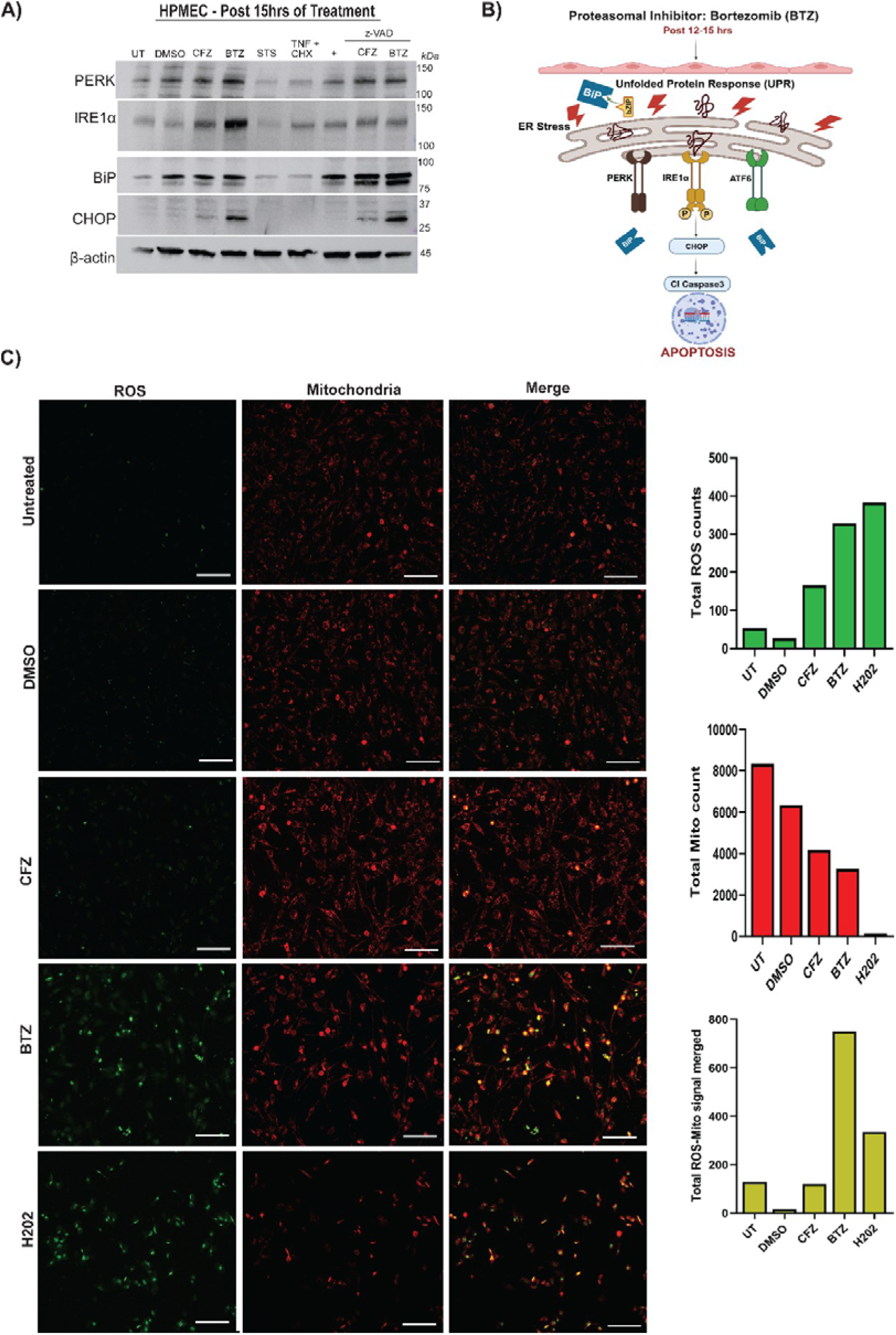
ER stress marker CHOP is activated via IRE1α in BTZ-treated HPMEC. (A) Representative immunoblot of ER stress markers in HPMECs treated with DMSO, CFZ (100 nM), BTZ (100 nM), STS (1 μM), TNF (25 ng/ml) + CHX (1 μM), zVAD (20-25 μM), and combinations of zVAD with CFZ, BTZ, or STS. Markers include PERK (140 kDa), IRE1α (130 kDa), CHOP (27 kDa), and BiP (78 kDa), with β-actin (45 kDa) as a loading control. (B) Schematic of ER stress pathway leading to CASP3 activation and apoptosis. (C) Fluorescence microscopy of HPMECs treated with CFZ (100 nM) or BTZ (100 nM) and controls (DMSO, H[O[500 μM). Cells were stained with CellROX Green (ROS) and MitoTracker Red (mitochondria). Representative images show ROS (green), mitochondria (red), and merged signals indicating ROS localization in mitochondria. Quantification of green (ROS), red (mitochondria), and punctate merged fluorescence intensity was performed (n = 1). Scale bar = 170 μm.

### BTZ Drives Mitochondrial Stress in HPMECs

As ER stress and mitochondrial stress occur together in stressed cells [53], we next examined mitochondrial stress (Figure 3C). A key indicator is the accumulation of reactive oxygen species (ROS) in this organelle. To assess mitochondrial stress, we used fluorescent dyes mitotracker (red) and anti-ROS dye (green) (Figure 3C). Untreated cells showed diffused red staining indicative of active mitochondria with some ROS production (Figure 3C first row). DMSO-treated cells showed a similar pattern (Figure 3C second row). We used hydrogen peroxide as a positive control for ROS production (Figure 3C fifth row) and detected an elevated green (ROS) signal along with reduced spread of Mitotracker (red) indicating mitochondrial shutdown. When we compared CFZ to BTZ, we found that BTZ drives visibly more ROS green signal and punctate Mitotracker red signal (Figure 3C third and fourth row).

Punctate Mitotracker signal is indicative of mitochondrial stress and loss of membrane potential, disrupting the even spread of Mitotracker. Notably, ROS accumulation localized in regions where the mitochondrial membrane potential and structural integrity were lost (Figure 3C, fourth row merged image). We quantified red fluorescence intensity, green fluorescence intensity, and their ratio, showing that BTZ- and hydrogen peroxide-treated cells exhibited higher values than CFZ-treated cells (Figure 3C right graphs) for the represented field of view. These findings confirm that BTZ induces intracellular stress in HPMECs by increasing mitochondrial ROS production, disrupting mitochondrial membrane integrity along with impacting the endoplasmic reticulum (ER). Together, these changes contribute to the enhanced cell death observed in Figure 1 and the activation of cell death markers Cl- CASP9, Cl-CASP8, Cl-CASP3 (Figure 2).

### BTZ Treatment of HPMECs is Not Inflammatory

In mouse models, BTZ-induced neuropathies depend on TNF signaling, with TNF pathway blockade reducing neuronal pain in patients [54]. These findings suggest that external TNF, an inflammatory cytokine and key mediator of inflammation and cell death, may contribute to the observed BTZ phenotype. Additionally, cell-extrinsic TNF engagement with TNF receptor 1 (TNFR1) is a primary driver of CASP8 cleavage [55]. To resolve whether BTZ-dependent CASP8 cleavage (Figure 2C) occurs via production of inflammatory cytokines including TNF, we performed a cytokine array using cell-free media from unmanipulated or treated (DMSO or CFZ or BTZ) cells (Figure 4A). Cells were treated for 24 hours, and supernatants from multiple replicates were pooled for analysis. The heatmap revealed that in each condition cells had a distinct inflammatory profile. Unmanipulated cells showed detectable levels of chemokines CCL-5, C5a, cytokines IL-8, G-CSF, chemokine-like cytokine MIF, and serine protease inhibitor Serpin E1. DMSO-treated cells showed a reduction in CCL-5, C5a, G-CSF, IL-8; an increase in MIF and SerpinE1 indicating that DMSO alters the threshold inflammatory signaling of these cells. Media from CFZ-treated cells displayed a decrease in CCL5, IL-8, MIF, and Serpin E1; increase in G- CSF when compared to DMSO. Media from BTZ-treated cells showed the lowest production of the released factors compared to DMSO or CFZ for all the chemokines, cytokines, and Serpin E1. These findings are not surprising, as BTZ is known to be anti-inflammatory due to its downregulation of NFκB transcriptional activity [56]. Thus, BTZ-induced death of HPMECs appears to be driven by cell-intrinsic factors, not extrinsic signals. TNF was not detected in the cytokine array. To further verify whether these cells produce TNF or not, we quantified TNF in media using ELISA. As CASP8 activation was not detected at 15 hours post-treatment but was detected by 24 hours post-treatment, we tested TNF production at both time points (Figure 4B upper and lower panel). No detectable levels of TNF were observed in any treatment group at the indicated time points (Figure 4B). TNF + cycloheximide control settings showed detectable TNF as expected. Therefore, CASP8 activation in BTZ-treated cells is not an effect of late TNF production from stressed cells but induced by the cell-intrinsic stress initiated by BTZ. Our data depicts that the mode of proteasomal inhibition (reversibly by BTZ or irreversibly by CFZ) significantly influences cell intrinsic and extrinsic factors that will likely influence organ injury as well as inflammation in patients.

**Figure 4:**
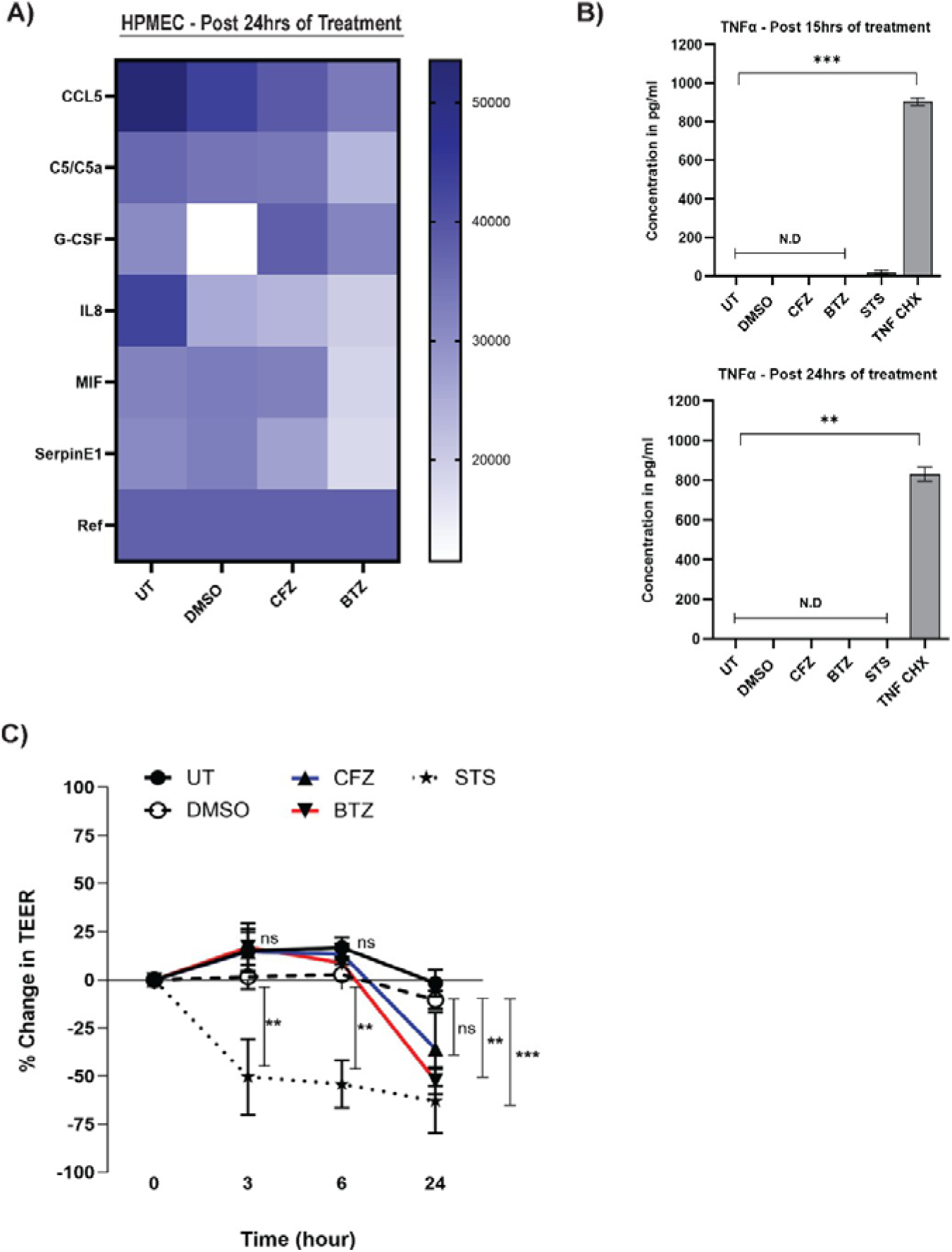
Effect of Bortezomib on Cell Extrinsic Signaling. (A) Heatmap of cytokine and chemokine levels in HPMECs after 24-hour treatments with DMSO, CFZ (100 nM), and BTZ (100 nM), measured using the Human Proteome Profiler Cytokine Array Kit. (B) TNF-α levels quantified by ELISA in HPMECs treated for 15–16 and 24 hours with DMSO, CFZ (100 nM), BTZ (100 nM), STS (1 μM), TNF (25 ng/ml) + CHX (1 μM). Statistical comparisons were performed using the Kruskal-Wallis test (12 hrs: **p < 0.01, 24 hrs: ***p < 0.001). (C) Trans endothelial electrical resistance (TEER) measurements of HPMEC monolayers treated with DMSO, CFZ (100 nM), BTZ (100 nM), and STS (1 μM). Percent change in TEER is shown, averaged from three independent experiments (± SEM). Statistical analysis: two-way ANOVA with multiple comparisons (DMSO vs. CFZ or BTZ, **p ≤ 0.01 at 24 hrs).

### BTZ confirms loss of cell-cell integrity of HPMECs

To assess the impact of BTZ-induced cell stress, death, and reduced inflammation on cellular integrity, we performed a Trans endothelial Electrical Resistance (TEER) assay. HPMEC monolayers modeling endothelial barrier were treated with 100 nM of BTZ, CFZ, DMSO, and STS (control). TEER measurements showed a decrease in resistance across all conditions over time (Figure 4C). At 24 hours post-treatment, BTZ, but not CFZ, significantly reduced resistance compared to DMSO. STS also caused a significant reduction throughout the time course. This confirms that BTZ increases HPMEC permeability [39], highlighting how BTZ disrupts endothelial homeostasis from within the cell while modulating essential cellular functions.

In summary, proteasomes are crucial for cellular homeostasis, a process cancer cells subvert for growth. Proteasomal inhibition, either natural (e.g., Alzheimer’s Disease) or during cancer chemotherapy (via CFZ or BTZ), leads to cell death and vascular integrity loss, which underlie BTZ-mediated cytotoxicity in vascular cells. These findings have implications for cancer therapy, clinical applications, and understanding diseases related to proteasomal inhibition.

## Discussion

BTZ has been a front-line chemotherapy drug for MM and other cancers for over two decades. There has been limited understanding of the mechanisms behind BTZ-induced adverse effects. Our study uncovers the mechanisms driving BTZ-induced cytotoxicity in HPMECs, which line pulmonary blood vessels. BTZ causes multi-organelle stress and activation of apoptotic pathways in HPMECs when directly compared to CFZ under equivalent conditions. Stress signals include increased expression of CHOP (Fig 3A), mitochondrial membrane compromise, and mitochondria-localized ROS (Fig 3C). In BTZ- treated HPMECs, the sequential IRE-1α and CHOP induction demonstrate that IRE-1, but not PERK pathway is a leading contributor to the ER stress. zVAD reduces IRE-1α levels in all conditions (Figure 3A) demonstrating that caspases contribute to PI-induced ER stress.

Our observations affirm prior studies implicating caspases upstream to ER stress [57]. The PERK levels remain comparable in zVAD-treated and untreated conditions (Figure 3A) suggesting that this pathway proceeds independent of caspases. As expected, zVAD restricts the appearance of CASP3 p17 fragment (Figure 2). PERK, IRE-1 or ATF-4-dependent ER stress responses are mediated by BiP, the primary sensor of ER stress. BiP binds to unfolded proteins and chaperones them to PERK, IRE-1 or ATF-4 [52]. We demonstrate that BiP is unchanged between CFZ or BTZ (Figure 3A). Thus, proteasomal inhibition by either compound activates the chaperone system. Intriguingly, pan-caspase inhibitor zVAD, increases intensity of BiP bands (Figure 3A) further indicating a potential caspase-dependent regulation in ER signaling upstream to the stress response. How BTZ involves ATF-4 response will require further analysis. Overall, inhibition of the proteasome irreversibly or reversibly, induces the ER chaperone system comparably in microvascular endothelial cells but has distinct outcomes for stress. What this means for other cell types lining the blood vessels (fibroblasts, smooth muscle cells) will require further investigation.

BTZ treatment induces both ER and mitochondrial stress in HPMECs simultaneously. Our work does not discern whether ER and mitochondrial stress pathways are parallel, sequential or synergistic to each other in this setting. CHOP-driven mitochondrial stress is a possibility that would implicate ER stress upstream of mitochondrial stress. CHOP, a developmentally regulated nuclear protein, triggers mitochondrial stress [58] and apoptosis by downregulating anti-apoptotic genes (Bcl2, Bcl-xl, Mcl-1), leading to the activation of Bim, Bax, and Bak.

CHOP can also collaborate with death receptors (DR-5, TRAIL) to activate CASP8 [59]. Another possibility is that BTZ directly affects mitochondrial proteases. A previous study demonstrated that BTZ binds to LonP1, a mitochondrial protease, but not carfilzomib [60]. LonP1 maintains mitochondrial health and prevents intrinsic apoptosis [61]. LonP1 inhibition also significantly activates the UPR and ER stress [62]. BTZ inhibits LonP1, triggering apoptosis through a mitochondria-mediated pathway. Therefore, we propose that in endothelial cells, BTZ stresses the ER and parallelly inhibits LonP1. This results in protein accumulation in mitochondria, proteotoxicity, and activation of the UPR in both the mitochondria and ER. In multiple myeloma cells, ER-mitochondrial stress nexus is a recognized effect of proteasomal inhibition and there is no resolution regarding the kinetics of events [49]. Any such effect of BTZ on regular cells has not been reported conclusively before ours. There are some indications that additional mitochondrial proteases may be targeted by BTZ. A previous study in human neurons demonstrated that BTZ binds to mitochondrial proteases cathepsin (Cat) G, catA, chymase, dipeptidyl peptidase II, and high-temperature requirement protein A2 (HtrA2)/Omi stress-regulated Endo protease. These findings argue for BTZ’s polypharmacologic effect and its ability to bind to multiple proteases. The authors conclude that HtrA2/Omi inhibition induces apoptosis in drug-treated cells [11]. In a contrasting study, authors demonstrated that BTZ does not target HtrA2/Omi [63]. There are more than 20 mitochondrial proteases, including HtrA2/Omi in the intermembrane space and LonP1 in the mitochondrial matrix, which are involved in stress responses at mitochondrial-associated membranes (MAMs). How mitochondrial proteases are targeted by BTZ and potentially underlie the observed cytotoxicity in vascular cells or other cell types require future-focused analyses. BTZ treatment recipients display various clinical adverse effects, including peripheral neuropathy, gastrointestinal toxicity, renal toxicity, thrombocytopenia, cardiovascular and severe pulmonary complications [64]. Recent findings have also implicated BTZ as an inducer of anti-proliferative, pathologic changes in lung arterial smooth muscle cells [65], further emphasizing the physiologic adverse effects of BTZ. While, our data do not address all these adverse effects individually, we bring to light important concepts of drug effects on vascular endothelial cells. Beyond, pulmonary complications, vascular damage may underlie the cardiovascular effects observed in BTZ recipients [66]. BTZ can be administered intravenously or subcutaneously, with studies suggesting that subcutaneous administration may be less toxic while maintaining similar efficacy [35]. Our data aligns with these findings and provides a potential mechanism about why intravenous administration is more toxic to patients. Previous research also have shown that BTZ affects cell cycle and permeability of vascular cells [67]. BTZ increases vascular permeability in HPMEC by reducing claudin-5, occludin, and VE- cadherin expression [39]. We also observe that BTZ drives significant endothelial cell permeability when compared to vehicle control DMSO (Figure 4C). Thus, it appears that BTZ has a multitude of effects on vasculature-cytotoxicity, changes in fundamental cellular function also changes in cellular inflammatory signals. The combination of cytotoxicity and loss of function potentially explains the multi-organ impaction observed in patients. Even though reduced inflammation is thought to be Vaso-protective [68], the reduced inflammatory signals from the HPMECs (Figure 4A) may be indicative of the reduced NFκB activity [78]. NFκB activity is critical in a healthy endothelial function and inflammatory response [56]. Incidentally, CFZ is thought to reduce ER stress and has anti-inflammatory as well as anti-thrombotic properties [69].

Future analyses will be necessary to show how irreversible blockage of the proteasome affects vascular health. In contrast to BTZ, CFZ appears to reduce hypertension [70]. Clinically, whether usage of CFZ before BTZ may reduce some of the adverse effects patients experience, needs to be investigated with utmost urgency. Our data also highlights the socio- economic aspects of multiple myeloma management [71]. BTZ is more affordable than CFZ, with generic versions available. Policymakers should consider updated data to create cost-effective strategies without compromising clinical outcomes or increasing adverse effects.

Although BTZ has proven effective, its toxicity necessitates the exploration of alternative regimens. Second-generation reversible proteasomal inhibitors, such as Ixazomib, prescribed as oral medication, should be investigated for toxicity and considered for broader clinical use [72]. In conclusion, there has been no prior mechanistic evidence of BTZ-induced ER-mitochondrial cytotoxicity. We provide novel insights into how proteasomal inhibition reversibly (by bortezomib) or irreversibly (by carfilzomib) impacts cancer (multiple myeloma) cells and non-cancer (pulmonary microvascular endothelial) cells.

## Resource Availability

Lead contact:

Further information and requests for resources and reagents should be directed to corresponding author and lead contact Dr. Pratyusha Mandal.

## Materials availability

All requests for materials should be directed to lead contact Dr. Pratyusha Mandal.

## Data and code availability

All data and any code are available through requests made to lead contact Dr. Pratyusha Mandal.

## Acknowledgments

This work was supported by the National Institutes of Health grant 1R15AI185729-01 (PM), National Institutes of Health grant SC2GM144168 (JGD), and Lehman College Startup Funds (PM). Additional support was provided by the Early Research Initiative (ERI) Pre- Dissertation Science Research Award 2024 from The Graduate Center, CUNY (PS) and NSF- LSAMP 2022 (AM). We thank Prof. Moira Sauane and Prof. Stephen Redenti (Lehman College, Department of CUNY) for cell culture reagents. We thank Prof. Edward Mocarski (Emory University), Hannah Wolf, and Jamie Price for providing feedback on the manuscript.

## Authors Contributions

Conceptualization: PS, PM

Methodology: PS, AM, JB, JGD, PM

Investigation: PS, AM, JB, PM

Visualization: PS, PM

Funding acquisition: PS, PM, JGD

Project administration: PM

Supervision: PM

Writing – original draft: PS, PM

Writing – review & editing: PS, AM, PM, JGD

## Declaration of interests

The authors declare that they have no competing interests.

## Supplemental information

## Materials and Methods

Figure S1. Bortezomib Induces Cell Membrane Permeability in HPMECs

Figure S2. Bortezomib Induces Caspase-Dependent Death

## Key resources

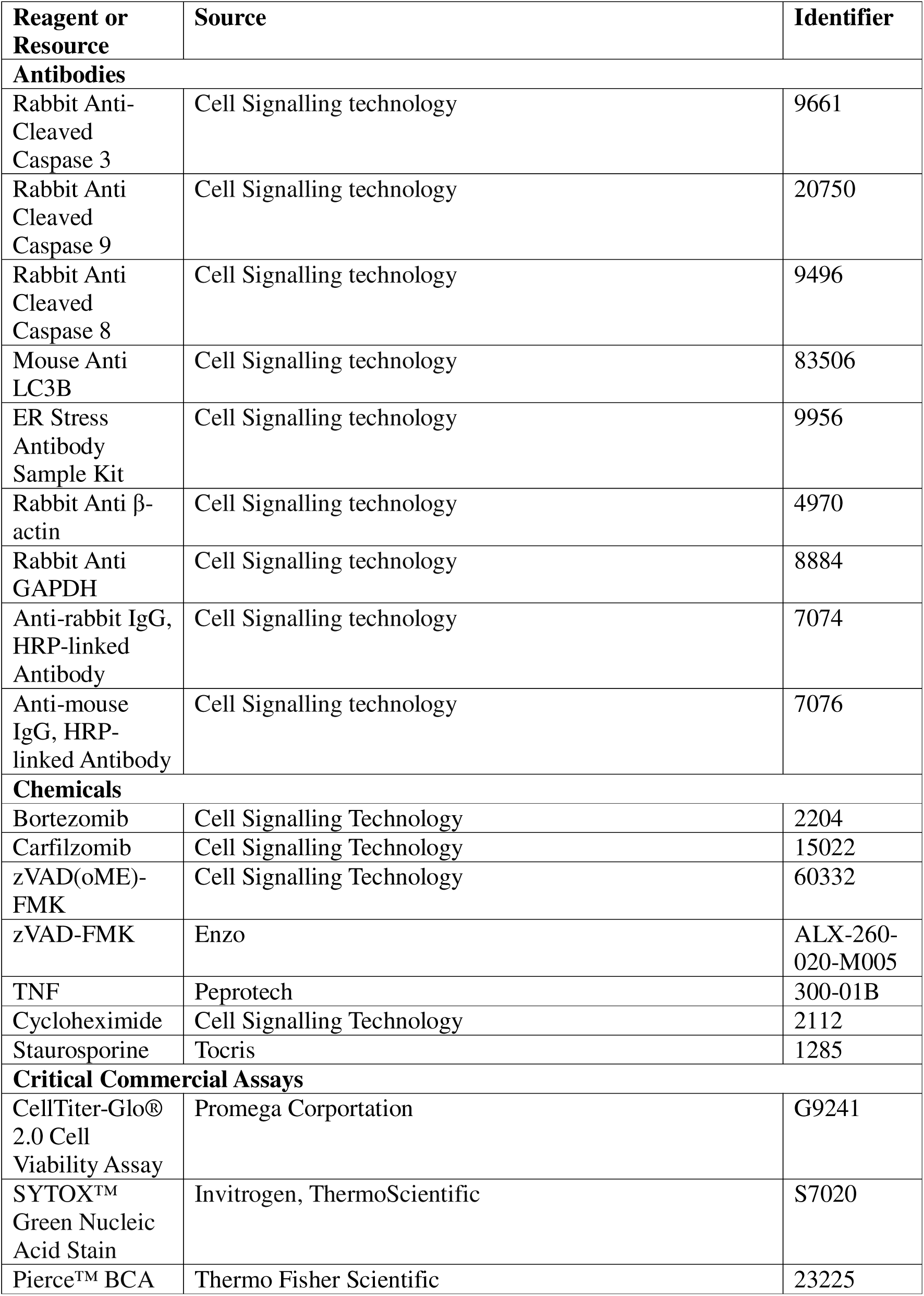
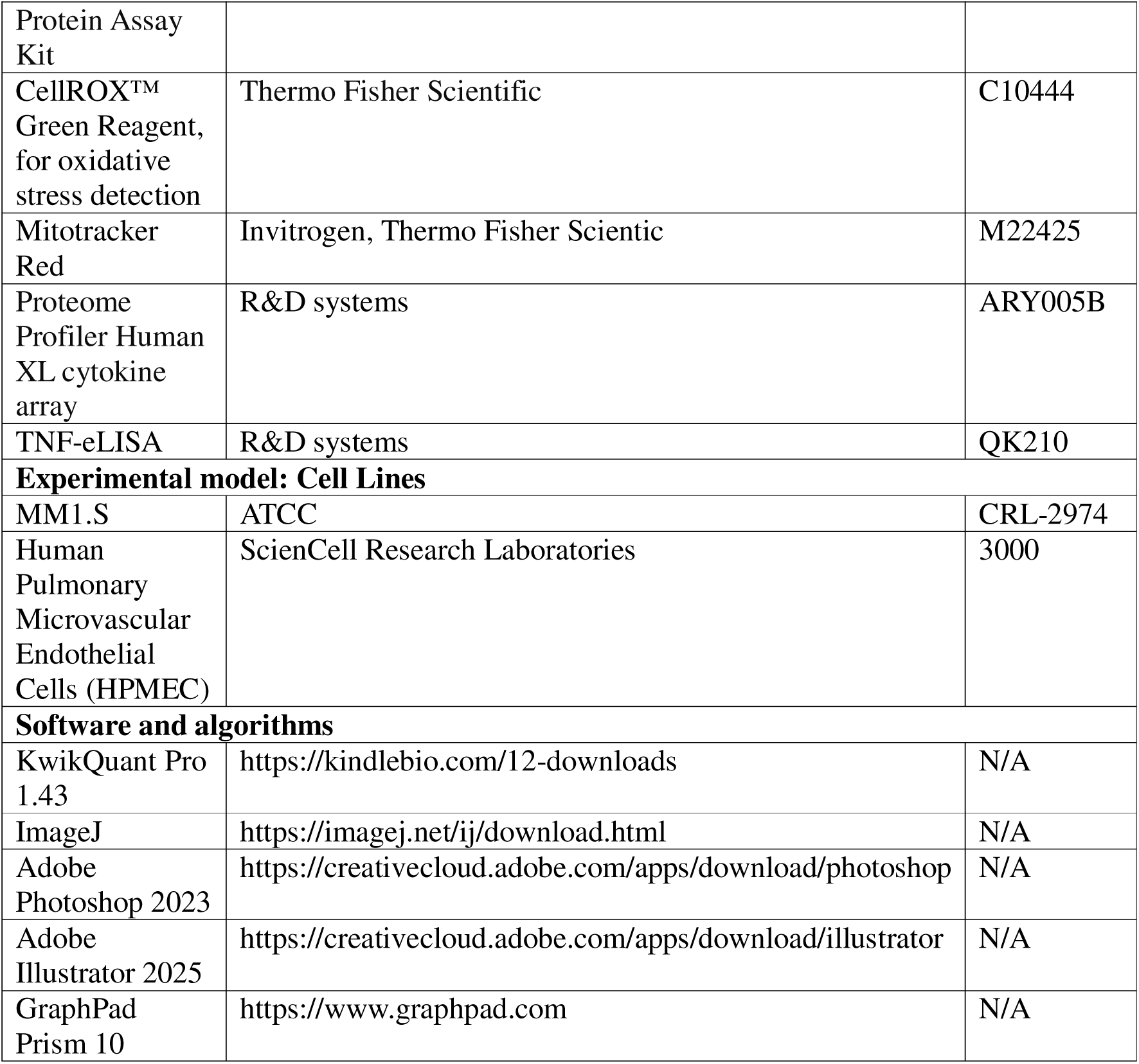

## Methods

### Cell Culture & Reagents

Human Pulmonary microvasculature endothelial cells (HPMECs) (ScienCell Research laboratories, Catalog 3000) were cultured in an Endothelial cell medium. (ScienCell, Catalog 1001) containing 5% of fetal bovine serum (FBS Catalog 0025), 1% of endothelial cell growth supplement (ECGS, Catalog 1052), and 1% of penicillin-streptomycin solution. (P/S, Catalog 0503) and Multiple Myeloma Cells (MM.1S) (ATCC, Catalog CRL-2974) were cultured in RPMI 1640 with 1% L-Glutamine (Corning, Catalog MT10104CV) supplemented with 10% Fetal Bovine serum (Gibco, Catalog A5256801) and 1 % antibiotic penicillin streptomycin strip (Corning, Catalog 30-002-CI), 1% L-Glutamine (Gibco, Catalog 25030081). All cells were maintained at 37^°^C in a humidified 5% CO2 incubator as per the ATCC guidelines. When reaching a cell density of 70% confluency all experiments were performed. FDA approved Chemotherapy drugs - Bortezomib (Cell Signaling Technology Catalog 2204), and Carfilzomib (Cell Signaling Technology Catalog 15022) at 100nM concentration were used for all the assays. Caspase inhibitors used were zVAD (oME)-FMK (Cell Signalling Catalog 60332) and zVAD-FMK (Enzo, Catalog ALX-260-020-M005). Cell stressors, TNF (Peprotech, Catalog 300-01B-1MG), Cycloheximide (Cell Signalling Technology, Catalog 2112), Staurosporine (Tocris, Catalog 1285) were used.

### Cell Viability Assay

Cell Viability was measured based on ATP detection using a CellTiter-Glo® 2.0 Cell Viability Assay kit (Promega Corporation, Catalog G9241) and Cell death was measured using SYTOX™ Green Nucleic Acid Stain (Invitrogen, Thermo Fisher Scientific Catalog S7020). The Viability assay based on ATP detection and SYTOX stain was performed as per the manufacturer’s instructions, seeded cells were treated with 100nM of Bortezomib and Carfilzomib for 24hrs for HPMEC and 12hrs for MM1.S. ATP luminescence was measured using Tecan (Model Spark 10M) at 550-570nm and for SYTOX fluorescent green signals of the cells were examined using a Echo Revolve Microscope (Model RVL2-K3).

### Immunoblot

For the extraction of proteins from the cells, the tissue culture flasks were scrapped using ice- cold TritonX-100 extraction Lysis buffer, pH 7.4 (ThermoScientific, Catalog J62289.AK) supplemented with Proteases inhibitor cocktail (Sigma-Roche, Catalog 11836170001) and Phosphatase inhibitor cocktail. (Roche, Catalog 4906837001). Cellular Content collected from the scrapped pellet was agitated in microcentrifuge tubes for 30 mins at 4^0^C, and centrifuged at 15000X g for 30 mins at 4^°^C. The Ultracentrifuged lysates were separated from the pellet and used for protein quantification and immunoblot. Protein Quantification was based on Bradford’s method using BSA as a control and performed according to the manufacturer’s instructions (Pierce™ BCA Protein Assay Kit, Thermo Fisher Scientific Catalog 23225). Samples were prepared using Laemmli SDS sample buffer (Thermoscientific, Catalog J61337.AD) was added to samples, and were boiled at 95°C for 5 minutes. Equivalent amounts of protein lysate (30-50ug) or equivalent volumes of cell lysates (20-30ul lysed supernatant) were loaded into SDS/PAGE gels (Mini-PROTEAN TGX, 4- 20%, Biorad, Catalog 456109) and separated proteins were transferred to PVDF membranes (Bio-Rad, Catalog 1620177). Membranes were then blocked with 5% BSA in TBS-T for 1hr at room temperature). Antibodies used were from Cell Signalling technology Anti- (Cleaved - Cl) Cl PARP (Catalog Asp214 D64610 5625), Cl CASP3 (Catalog D175 9661), Cl CASP9 (Catalog 20750), Cl CASP8 (Catalog 9496), LC3B (Catalog 83506), ER Stress markers - PERP, IRE1a, CHOP, BiP (ER Stress Kit Catalog 9956) β-actin (Catalog 4970), GAPDH (Catalog 8884) 1:1000 dilution in blocking buffer. Primary antibodies were incubated in cold temperature overnight. After overnight incubation, the membranes were washed three times with 1X TBST. Secondary antibodies used were Anti-Rabbit (Cell Signaling Technology, Catalog 7074) and Anti-Mouse (Cell Signaling Technology Catalog 7076 as per recommended dilution and incubated for 1hr at room temperature. Membranes were then washed with 1X TBST three times and developed using ECL western blotting substrate (Biorad, Catalog 1705061) and developed using KwikQuant Pro Imager (Model D1010).

### Detection of Reactive Oxygen Species and Mitochondria

Cellular ROS was detected using the CellROX™ Green Reagent, for oxidative stress detection (Thermo Fisher Scientific, Catalog C10444). Briefly, HPMECs were plated on 6 well plate and grown to 80% confluency. They were then treated with different drugs and treatment conditions for 15-16 hours. After treatment with drugs ROS was measured, 5uM CellROX green reagent was added to all the wells and incubated for 30mins at 37°C followed by 5mins PBS wash 3X, the cells were then stained with 50nM MitoTracker™ Dyes for Mitochondria Labeling (Invitrogen, Thermo Fisher Scientific M22425) for 30mins followed by 5mins PBS wash 3X, after all the washes cells were fixed with 15mins in 4% PFA. The fluorescent green and red signals of the cells were examined using Echo Revolve Microscope (Model RVL2-K3). Quantification of the fluorescent signals was performed using ImageJ software.

### Cytokine Array Analysis and ELISA

The Cell culture supernatant was collected from wells treated with drugs for 24hrs and was subjected to Cytokine analysis the samples were pooled for cytokine array (Proteome Profiler Human XL cytokine array, R&D systems Catalog ARY005B). The experiment was performed according to the manufacturer’s instructions. The membranes were stained using Streptavidin-HRP (Dilution, Company) and imaged using (BioRad ChemiDoc Imaging System). Analysis of the imaged membrane was done using Adobe Photoshop. For Elisa, Cytokines were quantified using ELISA kits for human TNF (R&D systems Catalog QK210) as per manufacturer’s protocol where cells were treated for 15-16hrs and 24hrs with drugs.

### Endothelial Permeability Assessment

Endothelial barrier integrity was assessed by measuring the trans-endothelial electrical resistance (TEER). HPMECs cells were seeded in a transwell cell culture inserts (Corning, Catalog 3470) in a 24-well plate. Cells were grown to confluency, then left untreated or treated with DMSO or CFZ or BTZ or Staurosporine. Cells were monitored over the time course 0, 3, 6, 24 hours. Resistance readings were taken using the EVOM3 TEER meter (World Precision instrument, Model EVOM3) as per the manufacturer’s instructions. TEER was calculated by multiplying the resistance by the surface area of the well and reported as a percent change.

### Statistical Analysis

Experiments are repeated at least three times with a minimum of two replicates in each set. Differences between means are determined by parametric, unpaired two-tailed Mann-Whitney (Figure 1), non-parametric Kruskal-Wallis test (Figure 4B) and two-way ANOVA with multiple comparisons (Figure 4C). P value <0.05 is considered statistically significant. All statistical analyses are performed using GraphPad Prism 10.

## Notes

### Competing Interest Statement

The authors have declared no competing interest.

