## Supplementary for "Cancer Drug Bortezomib, a Proteasomal Inhibitor, Triggers Cytotoxicity in Microvascular Endothelial Cells via Multi-Organelle Stress"

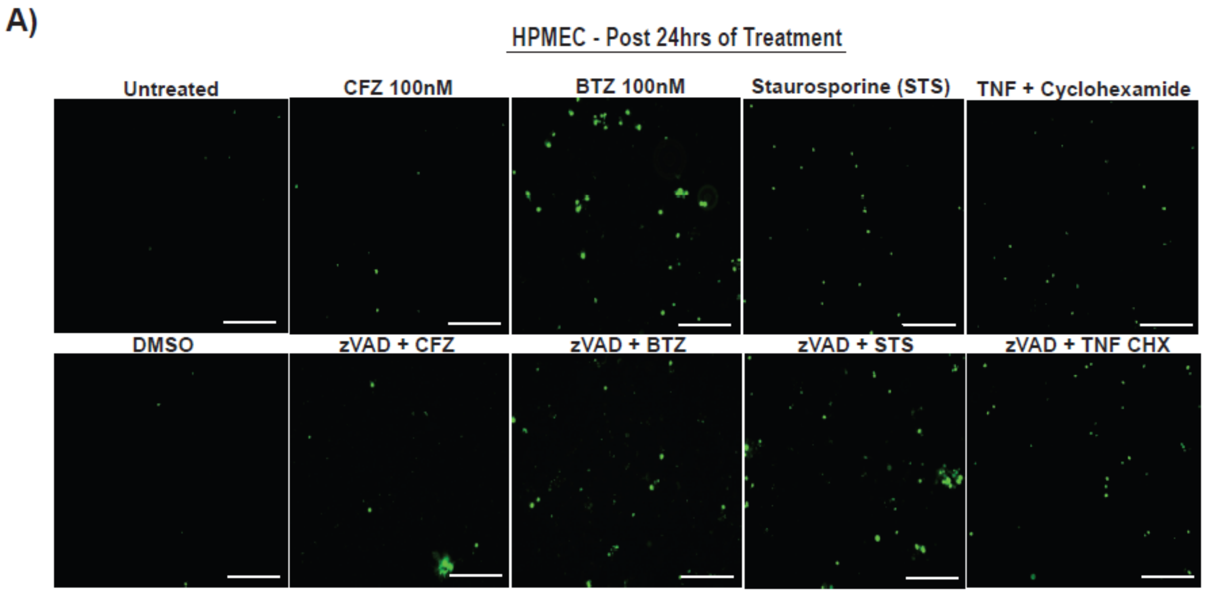


**Fig. S1. Bortezomib Induces Cell Membrane Permeability in HPMECs**

**(A)** Representative fluorescence microscopy images of HPMECs stained with SYTOX green (a cell impermeable dye) after 24 hours of treatment with DMSO, CFZ (100nM), BTZ (100Nm), STS (1 µM), TNF (25 ng/ml) plus Cycloheximide (CHX, 1µM), zVAD (20–25 µM), zVAD (20–25 µM) plus CFZ (100nM), zVAD (20–25 µM) plus BTZ (100nM), zVAD (20–25 µM) plus STS (1 µM), zVAD (20–25 µM) plus TNF (25 ng/ml) plus Cycloheximide (CHX 1µM). Scale bar = 170 µm. Images are representative of three independent experiments with n = 3 biological replicates in each experiment.

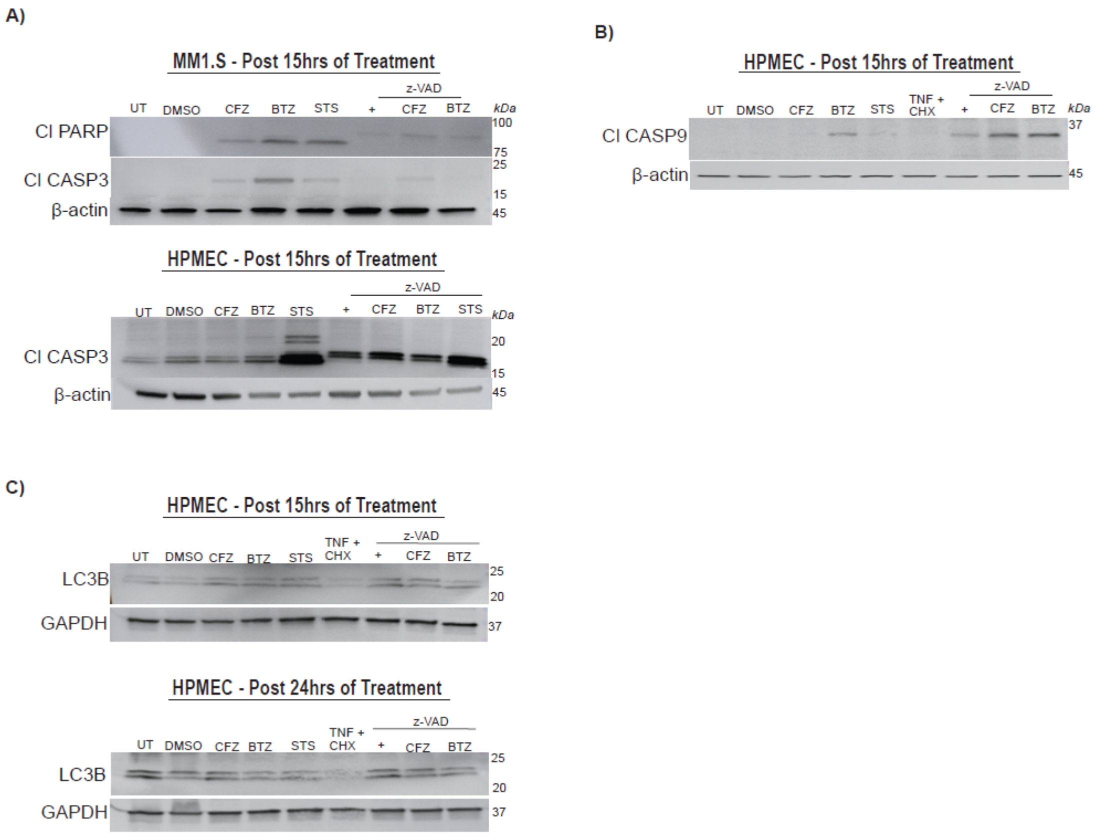


**Fig. S2. Bortezomib Induces Caspase-Dependent Death**

**(A-C)** Immunoblot analyses of the appearance of cell death markers at indicated times. Cells were treated with DMSO, CFZ (100 nM), BTZ (100 nM), STS 1 µM, TNF 25 ng/ml plus CHX (1µM), zVAD (20–25 µM), zVAD (20–25 µM) plus CFZ (100nM), zVAD (20–25 µM) plus BTZ (100nM), zVAD (20–25 µM) plus STS (1 µM) **(A)** MM1.S (upper panel) and HPMEC (lower panel) showing the expression of cleaved apoptosis markers: cleaved Caspase-3 (Cl-Casp3; 17, 19 kDa) and cleaved PARP (Cl-PARP; 89 kDa), with β-actin (45 kDa) as a loading control**. (B)** HPMEC showing intrinsic apoptosis marker cleaved Caspase-9 (Cl-Casp9; 35 kDa) and with β-actin (45 kDa) as a control. **(C)** HPMEC showing autophagy marker (LCB3 – 14,16kDa) and GAPDH (37 kDa) as a loading control.
